# The emergent mental operation of the learning brain

**DOI:** 10.64898/2026.06.03.729881

**Authors:** Tianyong Xu, Peng Tao, Jiali Mu, Xiao Han, Fan Liu, Peng Gao, Tian Xia, Zhu-Qing Gong, Hongjian He, Ting Xu, Luonan Chen, Feiyan Chen, Xi-Nian Zuo

## Abstract

How does the learning brain give rise to emergent mental operations that enable skill internalization and generalization? Using five-year longitudinal tracking of children acquiring abacus-based mental calculation, we reveal how sustained practice progressively reconfigures whole-brain states. Learning induces nonlinear transitions from stable baseline configurations through a transient exploratory phase toward restabilized associative brain networks. This hierarchical remodeling cascades from perceptual systems to action and default networks, mediated by adaptive neural variability and selective connectivity reweighting, with the salience-parietal memory network acting as a dynamic relay. These brain-state transitions accompany an ordered cognitive transfer cascade, emerging within arithmetic and extending to visuospatial and executive functions. Our findings are reproducible and provide a systems-level account of how ecological learning sculpts large-scale networks to enable complex skill acquisition.

## Main Text

Understanding how intelligence emerges from the interplay between innate constraints and experience-dependent plasticity is a central challenge for both cognitive neuroscience and artificial intelligence (*1*–*4*). Learning provides the primary substrate for this emergence and transforms our transient experiences into stable, transferable representational structures that expand the brain’s functional repertoire (*5, 6*). Although the synaptic and cellular mechanisms of plasticity are well characterized (*7, 8*), how such local changes accumulate to reshape whole-brain dynamics over extended periods of real-world learning remains largely uncharted (*9*–*11*). In particular, the spatiotemporal evolution of large-scale brain states supporting the transition from novice to expert has yet to be mapped at the systems level.

Theoretically, learning can be framed as an allostasis-oriented process through which the brain maintains functional stability by actively reorganizing its internal architecture in response to changing demands (*12*–*14*). Within this view, skill acquisition unfolds through an iterative cycle of exploration, selection, and refinement, driven by persistent mismatches between environmental challenges and available neural resources. Across learning, this adaptive trajectory is reflected structurally by an initial expansion followed by renormalization of neural resources (*11, 15*) and functionally by a shift from broadly integrated network configurations toward more specialized and efficient architectures (*16, 17*). Importantly, such plasticity unfolds in a coordinated, multiscale manner, encompassing synaptic modification, regional specialization, transient neural variability, and large-scale reorganization of network integration and segregation (*16, 18*–*23*). Accordingly, the internalization of expertise, defined as the transformation of effortful, externally scaffolded operations into automated, endogenous representations, is increasingly understood as a distributed network reorganization rather than a localized circuit refinement (*9, 24, 25*).

Despite these conceptual advances, empirical evidence for long-term, system-wide learning-induced dynamics remains limited. Much of what is currently known derives from short-term laboratory training or cross-sectional comparisons of experts, paradigms that offer experimental control but often lack ecological validity (*26, 27*). These limitations are evident in ongoing debates surrounding cognitive training, where narrowly defined interventions frequently yield inconsistent neural effects and weak behavioral transfer (*28*–*30*). More broadly, they reflect a growing recognition that simplified, highly constrained tasks may incompletely engage the brain’s intrinsic modes of organization. In contrast, naturalistic neuroscience has demonstrated that complex, context-rich experiences are essential for revealing how distributed brain networks operate and reorganize over extended timescales (*31, 32*). Together, these considerations underscore the need for longitudinal, ecologically grounded paradigms capable of capturing how prolonged real-world learning reshapes intrinsic brain network architecture to support the emergence of intelligent behavior.

Here, we address this gap using abacus-based mental operation (AMO) as a model of ecologically grounded, long-term skill acquisition (Fig. 1a). AMO involves the gradual internalization of a physical computational tool into a “virtual abacus in the brain,” rendering otherwise covert representational transformations explicitly tractable. Leveraging a longitudinal functional MRI (fMRI) cohort spanning up to five years (Fig. 1b), we delineate the neurocognitive trajectory from novice learning to advanced mastery. We show that the sustained AMO learning induces a nonlinear, progressive remodeling of brain states, orchestrated by coordinated interactions among visual, action, salience/parietal memory, and default networks. This network-level reconfiguration supports both efficient skill internalization and robust cognitive transfer beyond the acquired domain. By validation with an independent densely sampled dataset (Fig. 1c), our findings provide a systems-level account of how prolonged, ecologically embedded learning drives the emergence of intelligent behavior.

**Fig. 1.**
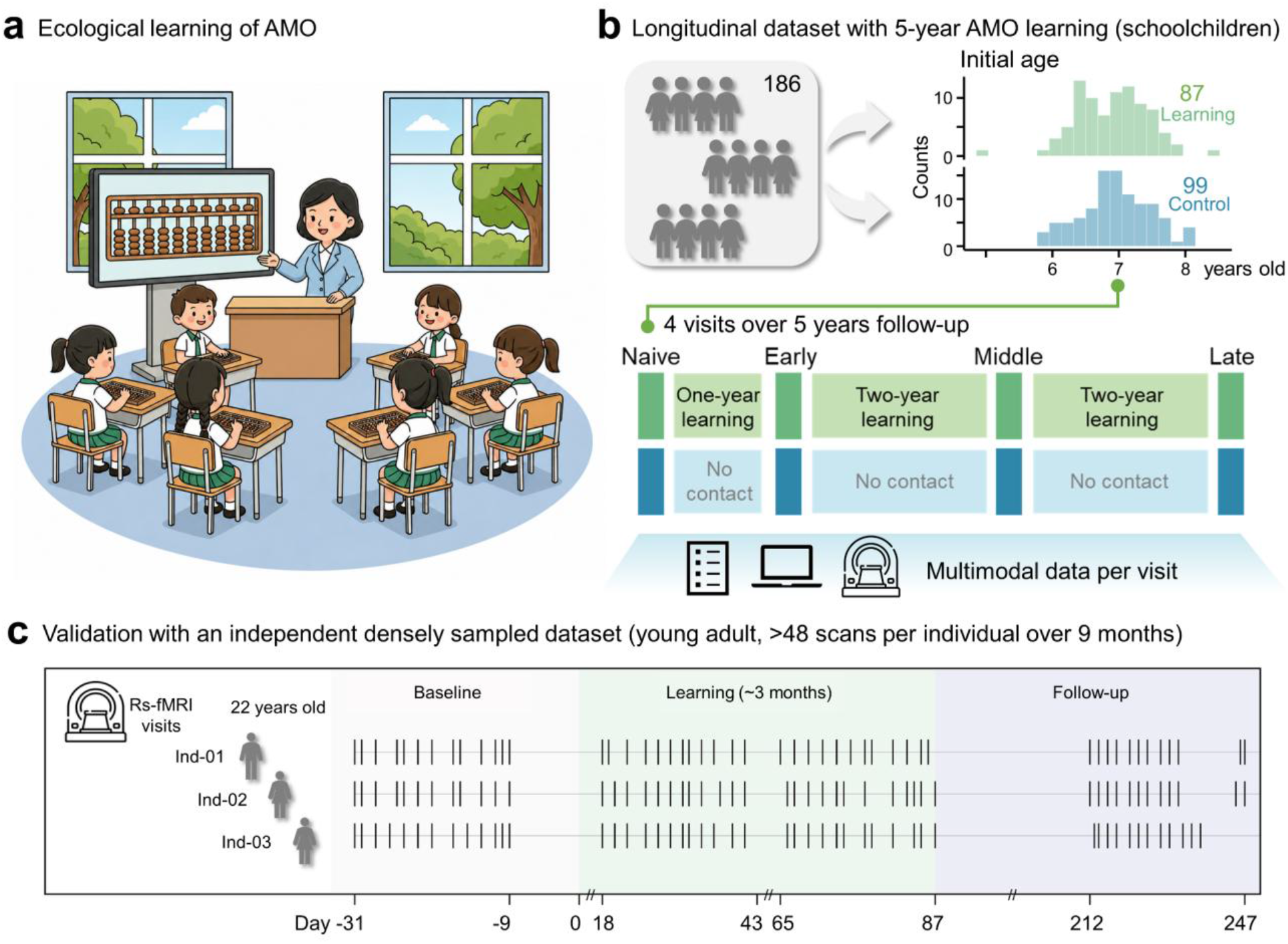
Longitudinal experimental design tracking the neurocognitive trajectory of AMO learning. **a**, Ecological learning of AMO. The incorporation of ecological learning of AMO into the school curriculum has the potential to facilitate extended and advanced learning. **b**, Primary longitudinal dataset with extended AMO learning. School-aged children were assigned to an AMO learning cohort or a non-intervention control group and followed over a five-year period. This design provides an ecologically grounded model of long-term skill acquisition spanning a critical developmental window. The learning and control groups exhibited comparable age and sex distributions throughout the study period, minimizing developmental confounds. **c**, Validation using an independent, densely sampled dataset. This comprised three young adults, with over 48 scans per individual, collected over a period of nine months.

## Results

To conceptualize system-level changes during long-term skill acquisition, we drew on the epigenetic landscape originally proposed by Conrad Waddington (*33, 34*) and modelled learning as a differentiation of trajectories in neural state space (Fig. 2a). Specifically, this differentiation manifests as a transition across distinct dynamical regimes induced by prolonged learning: the brain departs from an initial stable configuration and ultimately converges onto two different states.

**Fig. 2.**
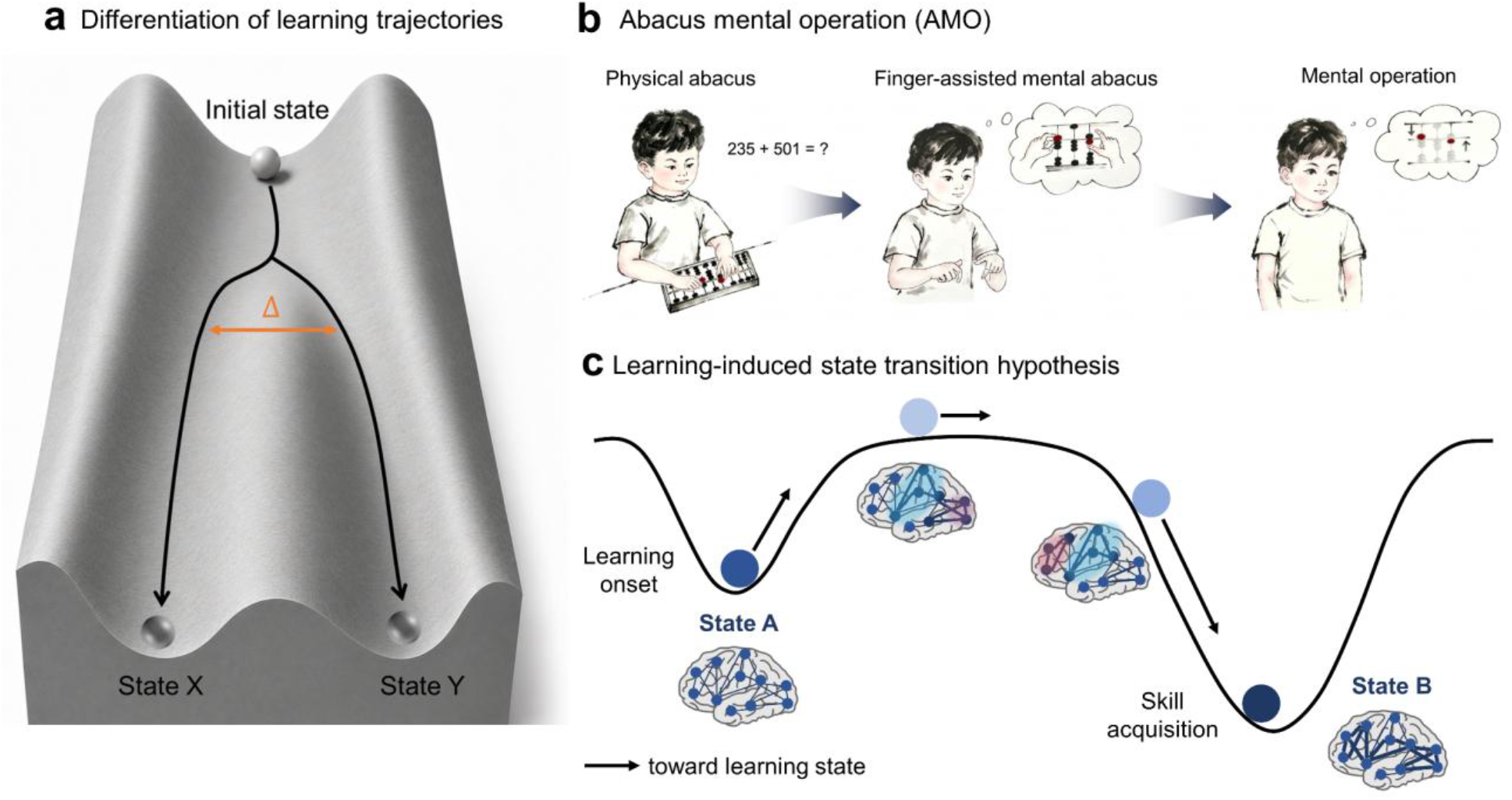
Learning trajectory differentiation on a neural landscape. **a**, Schematic illustration of learning as trajectory differentiation on a Waddington-inspired landscape, with transient destabilization enabling transitions between stable attractors. **b**, The AMO learning paradigm, progressing from physical abacus manipulation, to finger-assisted mental simulation, and ultimately to fully internalized mental calculation (‘virtual abacus’). **c**, A conceptual model of learning-driven brain state reconfiguration, illustrating the predicted progression from an initial stable configuration, through a non-stationary learning phase, to a reorganized, stabilized state.

Applied to AMO learning (Fig. 2b), this framework captures a characteristic progression in large-scale brain dynamics. Ongoing activity patterns are expected to transiently deviate from baseline stability during learning, reflecting heightened reconfiguration towards a non-stationary state, before stabilizing as newly acquired skills consolidate (Fig. 2c). Such transitions should be accompanied by coordinated changes in system-level reorganization rather than isolated regional effects.

To capture these dynamics, we applied a brain-state analysis informed by dynamic network bifurcation theory (*35*–*38*). This approach quantifies the mean nodal variability and inter-regional coupling in a coordinated fluctuation system, integrating these measures into a composite brain-state score that indexes both stationarity and global engagement during learning-related reconfiguration. Using this framework, we next tested whether prolonged AMO learning is associated with systematic, stage-dependent differentiation of brain-state trajectories and corresponding reorganization of large-scale network architecture.

### Nonlinear trajectory of brain-state reconfiguration during learning

Guided by this dynamical framework, we examined whether longitudinal AMO learning follows a differentiated trajectory of brain-state reconfiguration (Fig. S1). Brain-state analysis revealed a significant group × stage interaction in brain-state scores (*χ*^2^ (3) = 38.146, *P* < 0.001, ΔAIC = −32.15; Fig. 3a), indicating distinct reconfiguration dynamics between groups. Baseline configurations were comparable between the learning and control groups (*t*(66) = −1.855, *P* = 0.068, 95% CI [−0.467, 0.017]), whereas trajectories diverged following learning onset.

**Fig. 3.**
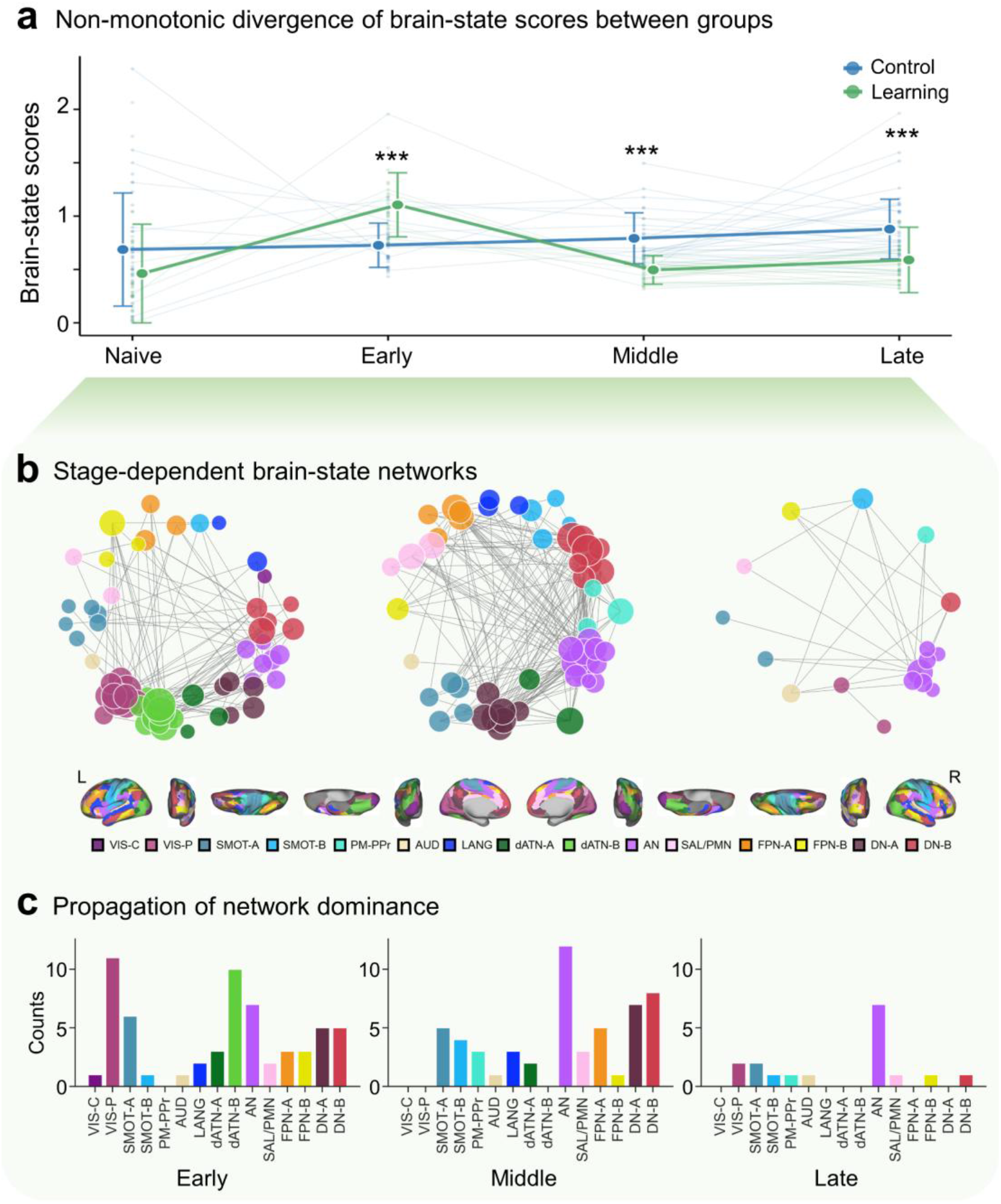
Nonlinear remodeling of brain states during long-term skill acquisition. **a**, Longitudinal trajectories of brain-state scores reveal a significant non-monotonic divergence between the learning and control groups (group × stage interaction, *P* < 0.001), characterized by comparable baseline configurations followed by a pronounced, learning-dependent reorganization. Error bars denote s.d. across participants. Note: \*\*\**P* < 0.001. **b**, Stage-dependent networks contributing to the group difference in brain-state scores, showing a progression from widespread early-stage networks to association networks at later stages. **c**, Spatiotemporal evolution of the dominant networks underlying the group difference. An early learning bias toward widespread reconfiguration in visual, dorsal attention and action networks progressively shifts toward higher-order associative networks, including the action and default networks, at later stages.

Specifically, in contrast to the stable profile observed in controls, the learning group exhibited a transient increase in state scores during the early learning stage (*t*(55) = 5.638, *P* < 0.001, 95% CI [0.244, 0.513]), followed by a progressive decrease across middle and late learning stages relative to controls (middle stage: *t*(56) = −5.868, *P* < 0.001, 95% CI [−0.397, −0.195]; late stage: *t*(61) = −3.902, *P* < 0.001, 95% CI [−0.437, −0.141]). Together, these results indicate that learning-related brain network reconfiguration follows a non-monotonic trajectory rather than a linear accumulation of network changes.

Mapping stage-dependent group differences in brain-state expression onto large-scale brain networks (*39, 40*) revealed a temporally ordered reconfiguration of network topography (Fig. 3b, S2 and S3). The early learning stage was characterized by widespread alterations in brain state reconfiguration, particularly within the visual, dorsal attention, and action networks. By the middle stage, differences within the visual and dorsal attention networks were no longer apparent, whereas alterations in the action and default networks became increasingly pronounced. Following prolonged learning, the action network emerged as the most stably reorganized system, suggesting a central role in sustaining long-term skill-related functional coordination (Fig. 3c).

### Progressive reconfiguration of functional network architecture

Focusing on the stage-dependent networks contributing to the group difference of brain-state scores, including visual, action, salience/parietal memory, and default systems (Fig. 4a), we examined learning-related reconfigurations of functional network architecture by quantifying the topological properties of the networks predominating at successive learning stages. within-network recruitment increased in the visual system during early learning, whereas recruitment within the action network declined at later stages (see Fig. 4b, top). In parallel, inter-network integration exhibited selective reconfiguration. Coupling between the action and salience/parietal memory networks was persistently reduced, whereas integration between the salience/parietal memory and default networks increased during the late learning stage.

**Fig. 4.**
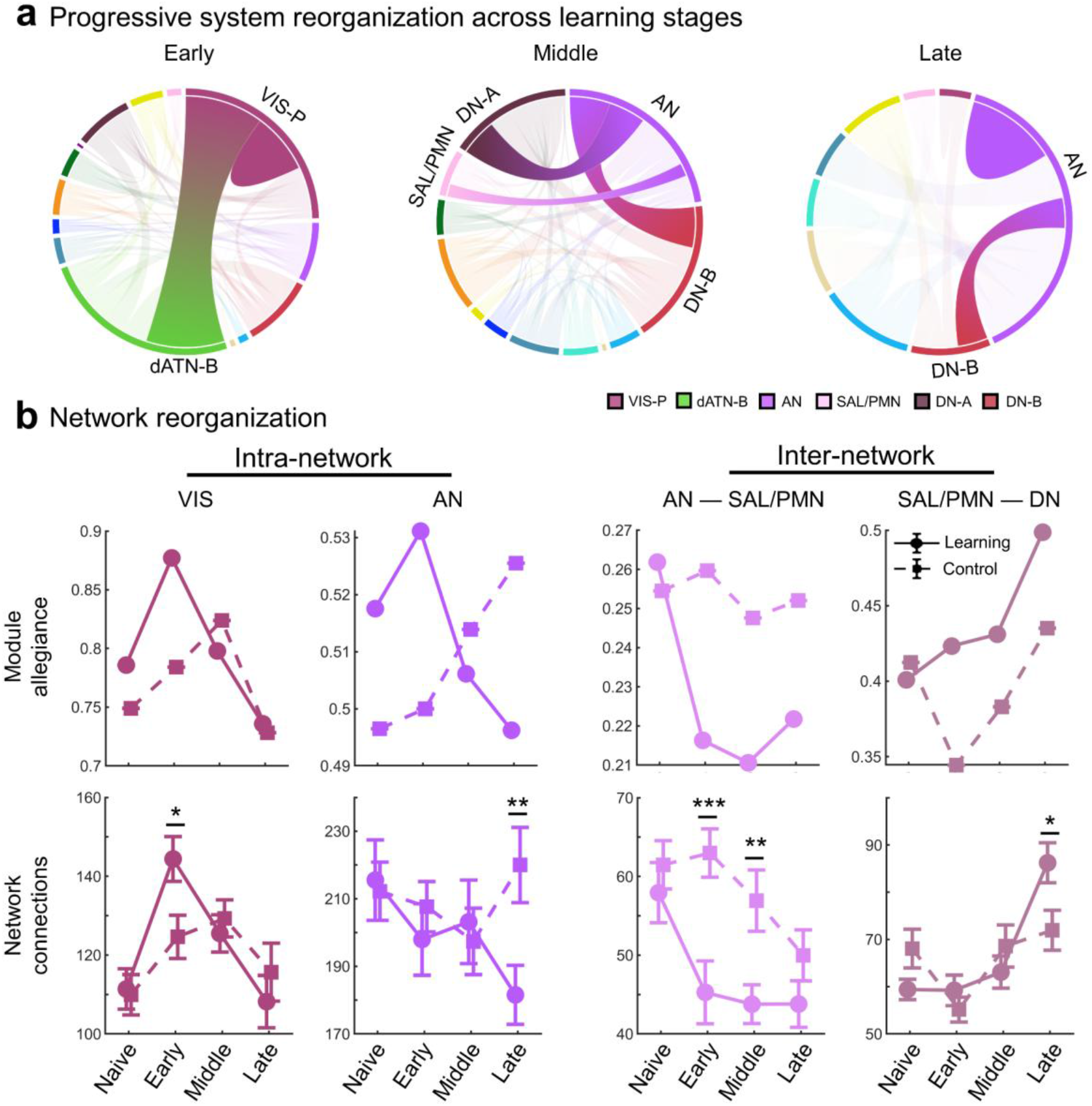
Progressive reconfiguration of large-scale brain networks during skill internalization. **a**, Conceptual summary of the stage-dependent trajectory of network reorganization involving visual, action, salience/parietal memory, and default networks. **b**, Reconfiguration of network topology. Top, modularity analysis showing changes in intra- and inter-network organization across learning stages. Bottom, corresponding changes in intra- and inter-network functional connectivity, revealing stage-specific reweighting of large-scale network interactions. Note: \*\*\**P* < 0.001, \*\**P* < 0.01, \**P* < 0.05.

Functional connectivity analyses converged on this topological shift (see Fig. 4b bottom). Intra-network connectivity within the visual system were selectively enhanced during early learning (*t*(54) = 2.361, *P* = 0.022, 95% CI [2.979, 36.526]), whereas connectivity within the action network decreased during the late stage (*t*(62) = −2.733, *P* = 0.008, 95% CI [−66.579, −10.330]). Inter-network coupling followed a similar progression: connectivity between the action and salience/parietal memory networks was reduced during early and middle stages (early: *t*(53) = −3.549, *P* < 0.001, 95% CI [−27.698, −7.696]; middle: *t*(55) = −2.912, *P* = 0.005, 95% CI [−22.216, −4.103]), whereas coupling between the default and salience/parietal memory networks increased significantly in the late stage (*t*(61) = 2.388, *P* = 0.020, 95% CI [2.321, 26.245]). Analyses of connectivity strength also revealed concordant patterns (Fig. S5). Collectively, these findings illustrate a progressive shift from visually dominated coupling toward stabilized integration among associative networks, including action, salience/parietal memory, and default systems.

### Ordered cascade of cognitive transfer mirrors neural reorganization

We next examined whether the large-scale neural reorganization during learning was accompanied by generalized cognitive benefits across behavioral domains. Cognitive transfer gains exhibited a graded pattern, with improvements diminishing across progressively more distal cognitive domains relative to the acquired AMO skill, from mathematics to executive domains (Fig. 5a).

**Fig. 5.**
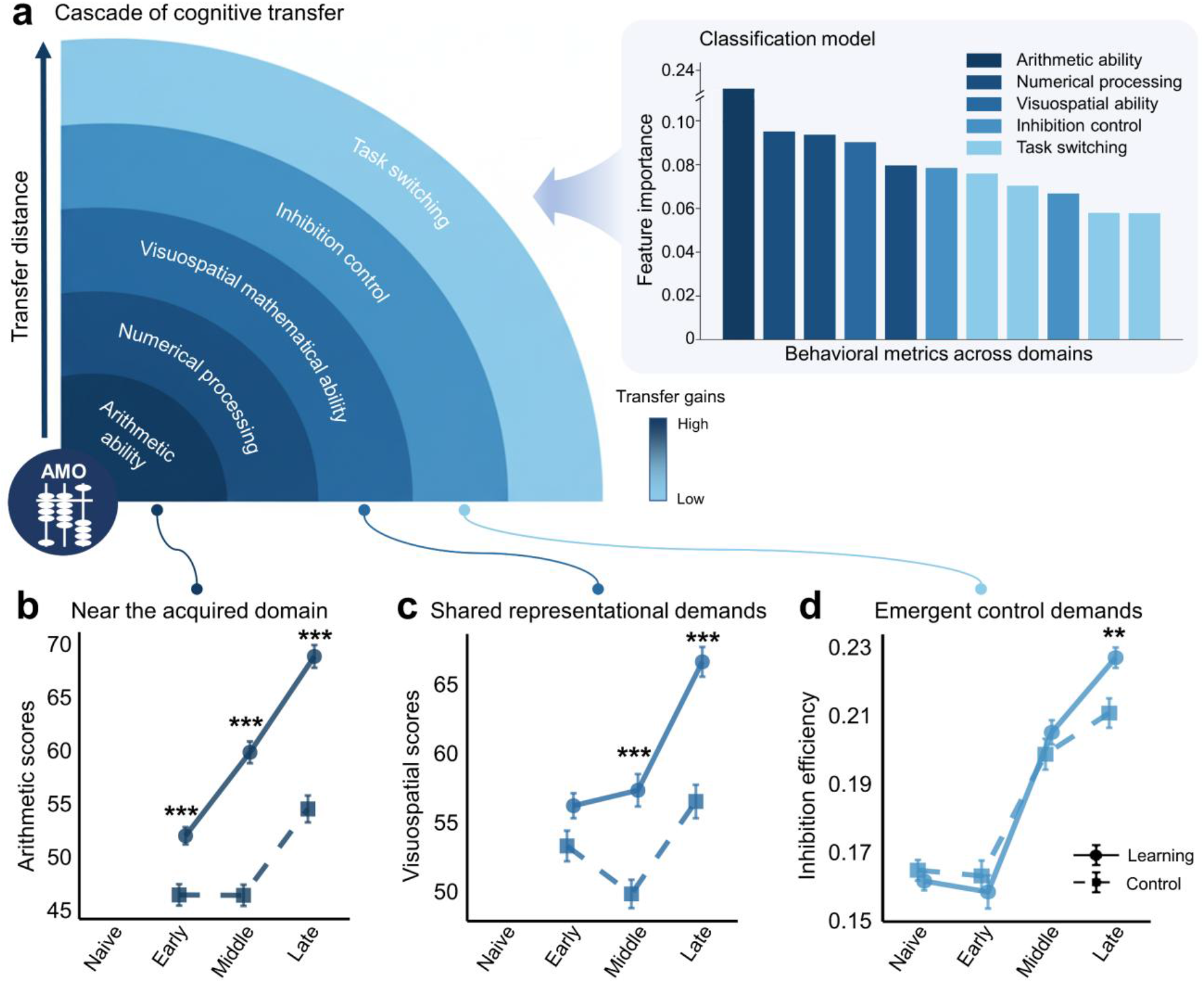
Sequential cascade of cognitive transfer tracks neural reorganization. **a**, Cognitive transfer effects across behavioral domains ranked by their distance from the acquired AMO skill. Transfer magnitude decreased progressively from arithmetic and numeric processing to visuospatial mathematical and executive functions. **b**, Arithmetic performance showing early emergence of near-transfer effects during learning. Error bars denote s.e.m. across participants. **c**, Visuospatial mathematical performance showing transfer effects emerging during the middle stage of learning. **d**, Inhibitory control performance showing transfer effects emerging during the late stage of learning, measured as inhibitory efficiency during a Go/No-go task (accuracy normalized by reaction time; higher values indicate better performance). Note: \*\*\**P* < 0.001, \*\**P* < 0.01.

This transfer hierarchy was supported by a random-forest classifier trained on post-learning behavioral measures, which reliably differentiated AMO participants from controls (AUC = 0.81; balanced F1 score = 0.72; accuracy = 0.74). Feature-importance analysis revealed a convergent ordering of predictive contributions, decreasing systematically from arithmetic ability to numeric processing, visuospatial mathematical skills, and executive functions (Fig. 5a, right).

Consistent with this behavioral hierarchy, performance gains emerged earliest in arithmetic ability (Fig. 5b; group × stage interaction, *χ*^2^ (2) = 51.115, *P* < 0.001, ΔAIC = −47.1; significant group difference in all learning stages, all *P*s < 0.001), subsequently extended to visuospatial mathematical processing (Fig. 5c; *χ*^2^ (2) = 25.15, *P* < 0.001, ΔAIC = −21.1; significant group difference in middle and late stages, all *P*s < 0.001), and later appeared in executive control, specifically inhibitory function (Fig. 5d; *χ*^2^ (3) = 17.505, *P* < 0.001, ΔAIC = −11.5; significant group difference in late stage, *P* = 0.009). Collectively, these results indicate that prolonged learning induces a progressive expansion of cognitive transfer from task-proximal abilities toward more general cognitive functions, paralleling the stage-dependent stabilization of large-scale brain network organization.

### Independent replication in a densely sampled longitudinal cohort

To assess the reproducibility of the observed learning-related brain-state dynamics, we analyzed an independent validation cohort using a densely sampled longitudinal design. Despite substantial inter-individual variability, the principal trajectory identified in the primary cohort was preserved. This dataset comprised three learning stages: a baseline stage characterized by number-based mental arithmetic (NMA), an intermediate learning stage following initial acquisition of AMO, and a follow-up stage corresponding to proficient AMO performance. By comparing resting-state and task-based fMRI across stages, we examined whether learning-related shifts in brain states and network organization could be independently replicated.

Brain-state configurations progressively diverged from baseline through the learning stage and into the follow-up period (Fig. 6). During resting state, the transition from baseline NMA to proficient AMO was primarily associated with reorganization of sensorimotor systems (Fig. 6a). Brain-state scores differed significantly across stages (all *P*s < 0.05), a pattern that was also evident at the individual level (Fig. S6a).

**Fig. 6.**
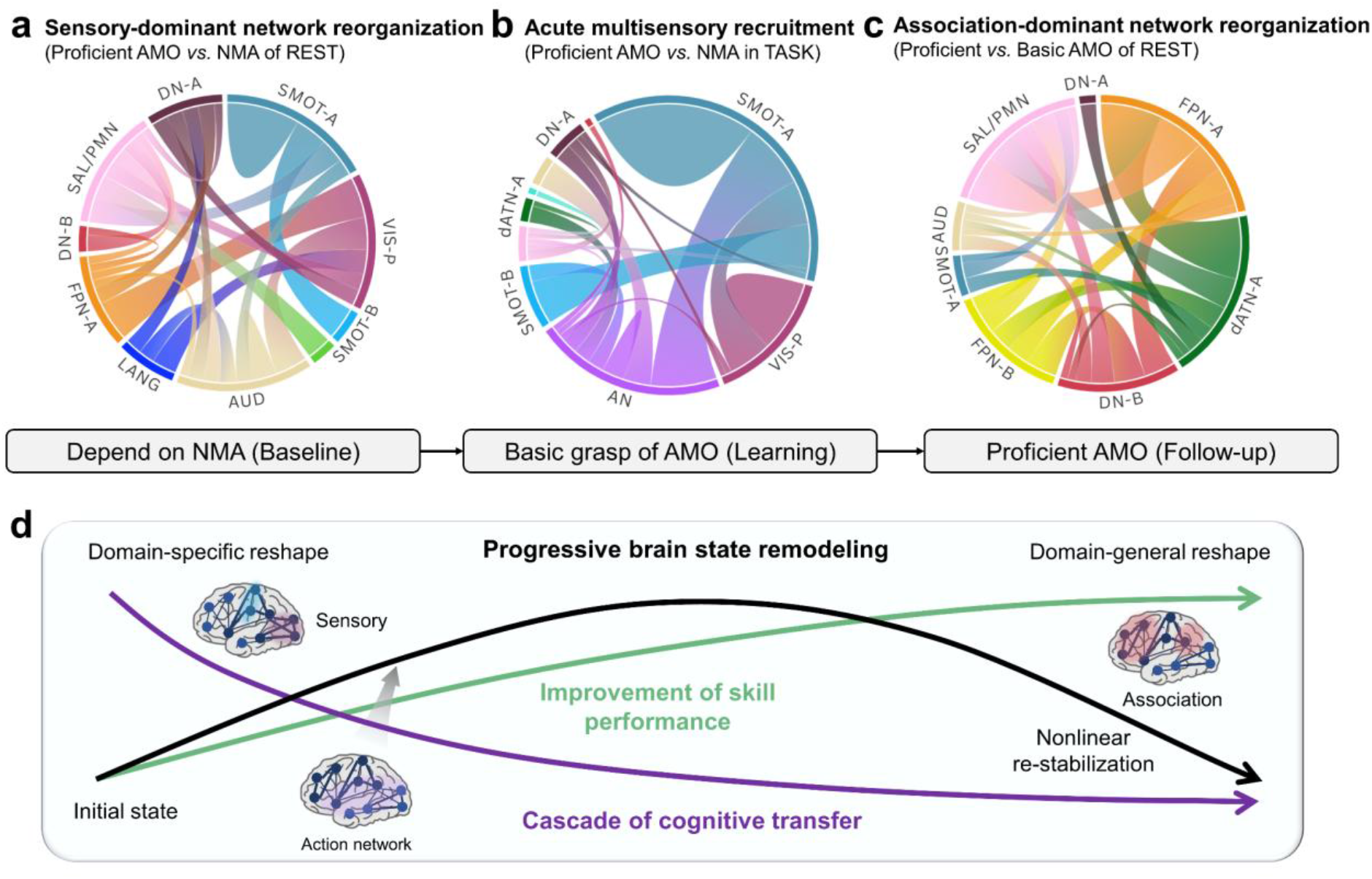
Brain-state dynamics in an independent longitudinal abacus learning cohort. **a**, Resting-state brain-state reorganization across learning stages. Differences between baseline number-based mental arithmetic (NMA) and follow-up proficient AMO stages were predominantly associated with sensorimotor-network contributions. **b**, Task-related brain-state reorganization during mental arithmetic. In addition to persistent sensorimotor involvement, the action network emerged as a major contributor to stage-dependent brain-state differences during AMO performance. **c**, Late-stage brain-state transitions. Significant differences between the learning and follow-up stages were accompanied by a redistribution of dominant network contributions toward higher-order association systems. **d**, Conceptual synthesis of learning-related brain-state dynamics. Long-term skill acquisition is associated with a differentiated trajectory of brain-state reorganization, progressive improvement in task performance, and the sequential emergence of cognitive transfer across domains increasingly distant from the trained skill.

Task-based fMRI acquired during mental arithmetic revealed a complementary expression of this reorganization. Whereas sensorimotor networks continued to contribute prominently throughout learning, the action network emerged as a major contributor during mental operations (Fig. 6b; Fig. S6b, all *P*s < 0.001). Notably, the relative contribution of the action network increased with AMO proficiency, paralleling the behavioral transition from externally guided calculation to internally generated computation.

A further transition between the basic and proficient AMO stages was observed in resting-state networks (all *P*s < 0.001, Fig. S6c). This later phase was characterized by a redistribution of dominant network contributions from sensorimotor systems toward higher-order association cortices (Fig. 6c), indicating continued reorganization beyond initial skill acquisition.

Together, findings from this densely sampled longitudinal cohort independently replicate the stage-dependent reconfiguration of brain states observed in the primary cohort. Across learning stages, brain-state organization shifted from sensorimotor-dominated configurations toward greater involvement of association networks, while task-state analyses highlighted a persistent contribution of the action network during skilled performance. The convergence of resting-state and task-state evidence across independent datasets suggests that progressive brain-state remodeling represents a reproducible feature of long-term skill acquisition rather than a cohort-specific effect.

## Discussion

Long-term learning is widely assumed to reshape brain organization, yet how this transformation unfolds across large-scale neural systems has remained poorly understood. By combining a five-year longitudinal learning cohort with an independent densely sampled validation dataset, we delineate the nonlinear trajectory of brain-state reorganization accompanying sustained skill acquisition. Rather than emerging through progressive strengthening of isolated task-specific regions, expertise was associated with differentiation of brain-state trajectories, characterized by an initial period of heightened reconfiguration followed by convergence toward stabilized network architectures. This transition was accompanied by an ordered reorganization of large-scale brain networks and a sequential expansion of cognitive transfer across behavioral domains. Together, these findings suggest that prolonged learning is best understood not as the accumulation of local neural changes, but as a non-monotonic process of system-level state-space reorganization through which experience becomes internalized and generalized.

Brain-state analyses revealed that learning followed a differentiated, non-monotonic trajectory through neural state space. An initially stable baseline configuration gave way to a transient phase of elevated reconfiguration during early learning, before converging toward a restabilized configuration at later stages. This pattern is consistent with an allostasis-oriented view of neural adaptation, in which functional stability is maintained not by resisting change but by dynamically reorganizing large-scale neural systems in response to evolving computational demands (*14, 41*). Within this framework, learning-induced plasticity serves not only to improve task performance, but also to recalibrate the balance of distributed neural systems during prolonged adaptation. More broadly, these findings suggest that learning may proceed through transient departures from established operating regimes, allowing the system to explore alternative configurations before stabilizing around more efficient solutions.

A central feature of this process is adaptive neural variability. Early learning was characterized by increased variability within visual systems, whereas later stages exhibited reduced variability among higher-order associative networks, including the action and default systems. Neural variability has been proposed to balance trait-like stability with task-dependent flexibility across cortical hierarchies and development (*42*–*44*). Elevated variability may facilitate exploration by enabling the system to sample alternative network configurations (*45, 46*), whereas subsequent reductions likely reflect consolidation of efficient architectures and improved energetic efficiency (*47*). From this perspective, transient destabilization is not a by-product of learning but a functional component of adaptation, enabling transitions between distinct organizational states.

Importantly, learning-related reorganization unfolded as a hierarchical differentiation of large-scale brain systems. Early learning was characterized by widespread reconfiguration centered on perceptual networks, particularly the visual system, whereas later stages increasingly involved associative systems, including the action and default networks. Concurrently, network organization shifted from broad within-network adjustments toward selective reweighting of inter-network interactions. This progression is consistent with theories proposing that expertise emerges through a redistribution of processing demands from externally guided perceptual operations toward internally coordinated representations. The accompanying increase in network segregation, a hallmark of efficient and specialized information processing (*16, 17, 48*). suggesting that prolonged learning promotes not only stronger representations but also more economical large-scale network organization. Convergent evidence from motor and cognitive learning studies indicates that prolonged practice is accompanied by systematic reconfiguration of large-scale brain networks, including redistribution of functional engagement across sensory, motor, and higher-order association systems (*9, 16, 17*). Together with the present findings, these observations raise the possibility that expertise emerges through a common process of learning hierarchical differentiation, whereby neural activity progressively transitions from broadly distributed and exploratory configurations toward increasingly specialized and coordinated network states.

Within this progression, the action network exhibited a learning-specific pattern of reorganization that diverged from normative development. Although functional connectivity strength increased comparably in both groups, suggesting preserved maturational growth (*49*), within-network recruitment declined selectively in the AMO learning group. This dissociation indicates that expertise is not necessarily associated with strengthened local cohesion but may instead arise through redistribution of functional engagement toward more selective and efficient integration. Sustained learning may therefore accelerate the transition of action-related systems toward mature operating regimes that support flexible coupling with higher-order associative networks. Crucially, because developmental increases in connectivity strength were comparable across groups, these effects are unlikely to be explained solely by maturation, highlighting a distinction between developmental growth and experience-dependent reorganization.

At the mesoscale, the salience/parietal memory network may emerge as a potential mediator of this hierarchical transition. As learning progressed, coupling between the action network and the salience/parietal memory network decreased, whereas interactions between the salience/parietal memory and default networks increased. Recently characterized as a functionally integrated system, this network occupies a strategic position for coordinating interactions between externally driven and internally oriented processing streams (*50*). Its established roles in salience detection, memory retrieval, and perceptual–response integration (*51*–*54*) suggest that it may facilitate the gradual transition from effortful computation to internally simulated and automated strategies. Through this mechanism, learning-related representations may become progressively detached from externally guided operations while remaining accessible through stabilized associative architectures.

Behaviorally, cognitive transfer emerged as an ordered cascade extending from arithmetic performance to visuospatial mathematical processing and subsequently to executive control. This sequence closely paralleled the progressive reorganization of large-scale brain networks and is consistent with models proposing that overlap in neural substrates constrains both the direction and extent of transfer (*55*). Evidence from an independent abacus-learning cohort further supports this interpretation, demonstrating that early learning gains are mediated primarily by domain-specific numerical processes, whereas later benefits increasingly depend on domain-general cognitive mechanisms (*56*). Notably, transfer effects emerged after prolonged learning, suggesting that generalized cognitive benefits may depend on system-level reorganization processes that unfold over timescales substantially longer than those typically examined in laboratory-based interventions. The ecological complexity of AMO learning—which integrates visuospatial imagery, working memory, attention, and motor planning into a unified and progressively internalized strategy—may have been particularly effective in promoting such distributed reorganization (*57*–*59*).

Several limitations should be considered. First, although the longitudinal design provides strong leverage for characterizing developmental learning trajectories, the primary cohort involved a specific form of cognitive expertise, and therefore the universality of the observed dynamics remains to be established in future research. Second, the densely sampled validation dataset consisted of a small number of individuals, although its intensive within-subject sampling provided complementary evidence for reproducibility. Third, while the present analyses characterize large-scale functional reorganization, future work integrating structural connectivity, electrophysiological recordings, and computational modeling will be necessary to clarify the mechanisms through which learning-related transitions between brain states are implemented.

In summary, prolonged learning was associated with a differentiated trajectory of brain-state reorganization, progressing from transient exploratory configurations toward stabilized and specialized network architectures. Across independent longitudinal datasets, this transition was accompanied by progressive reconfiguration of large-scale brain networks and an ordered expansion of cognitive transfer from domain-specific to increasingly domain-general functions. Beyond abacus learning, these findings suggest that the emergence of expertise may be governed by a common systems-level principle: prolonged learning differentiates trajectories through neural state space, enabling the transformation of externally guided operations into internally stabilized and transferable representations.

## ACKNOWLEDGEMENTS

The authors thank all the children, parents, and teachers who participated in the study.

## Author contributions

T.-Y.X., P.T., T.X. (Ting Xu), L.C., F.C., and X.-N.Z. conceived the study; T.-Y.X. and P.T. formulated the methodology; F.C. designed the experimental architecture and established the core datasets; J.M., X.H., and F.L. collected and curated the dataset; T.-Y.X. conducted the primary data analysis and visualization, with P.T. contributing to the analytical framework; J.M., P.G., Z.-Q.G., and T.X. (Tian Xia) provided neuroimaging preprocessing support; H.H. provided technical guidance on MRI protocols; T.-Y.X. wrote the original manuscript with input from P.T.; L.C., F.C., and X.-N.Z. supervised the study; All authors reviewed and edited the manuscript and approved the final version.

## Data, code, and materials availability

The CATP dataset is partially accessible at https://doi.org/10.1038/s41597-025-06500-9. The PEARL dataset, described in a neuro-resource article currently under review, will be made openly available upon publication. Requests for the datasets can be addressed to F.C.

