## Supplemental Materials for "The emergent mental operation of the learning brain"

**The PDF file includes:**

Materials and Methods
Supplementary Text
Figs. S1 to S6
Tables S1 to S2
References

**Other Supplementary Materials for this manuscript include the following:**

None

### Materials and Methods

#### Datasets and study design

We analyzed two complementary longitudinal datasets to characterize large-scale brain-state dynamics during skill acquisition: a population-level primary longitudinal cohort and an independent dense-sampling validation cohort. The primary cohort was used for all group-level statistical inference, whereas the dense-sampling cohort served to independently validate the temporal expression and individual specificity of the identified brain-state dynamics.

*Dataset 1 (Primary longitudinal cohort)*. The primary dataset was drawn from the Chinese Abacus Training Project (CATP), comprising 186 children (6–8 years at baseline), including a learning group ($n$ = 87) receiving five years of abacus-based mental operation (AMO) instruction and an age- and sex-matched control group ($n$ = 99, Table S1). A subset of 135 participants underwent longitudinal resting-state fMRI (287 scans in total, Fig. S1), enabling characterization of brain-state trajectories across extended learning. Cognitive assessments spanning mathematical ability and executive function were administered at multiple stages of the study (see Supplementary Text).

*Dataset 2 (Dense-sampling validation cohort)*. To validate the mechanistic features of the brain-state dynamics observed in the primary cohort, we analyzed an independent dense-sampling dataset (Precision Abacus-based Representation Learning, PEARL) comprising three adult participants undergoing intensive AMO learning with repeated fMRI scanning (over 48 scans) over nine months. This dataset was analyzed using the same brain-state framework but was not used for statistical inference or parameter estimation.

All participants had normal or corrected-to-normal vision and reported no history of neurological or psychiatric disorders. Written informed consent was obtained from all participants or their legal guardians. The study protocol was approved by the Ethics Committee of Zhejiang University.

#### MRI acquisition and preprocessing

In the primary cohort, resting-state fMRI data were acquired using a conventional single-echo EPI sequence and preprocessed using a standard volumetric pipeline including motion correction, nuisance regression, normalization to MNI space, and temporal band-pass filtering. In the validation cohort, resting-state and task-state fMRI data were acquired using a multi-echo multiband sequence and preprocessed using a dedicated multi-echo pipeline incorporating ME-ICA denoising and surface-based registration. Detailed acquisition parameters and preprocessing steps for each dataset are provided in the Supplementary Methods. MRI acquisition and preprocessing differed slightly between the primary and validation datasets, reflecting their distinct study designs. All analyses were conducted within each dataset using internally consistent preprocessing pipelines.

#### Brain-state analysis

To characterize large-scale functional reorganization during prolonged learning, we employed a brain-state analysis framework to compute a composite brain-state score for each individual. This framework is conceptually grounded in dynamic network bifurcation theory (*1*, *2*) and quantifies coordinated changes in regional signal variability and inter-regional functional coupling within a learning-relevant subnetwork (*3*). Higher brain-state scores indicate a relative departure from stationary brain dynamics toward systematically engaged, non-stationary states during learning. Brain-state scores are interpreted as relative indices within each dataset and are not intended for direct numerical comparison across datasets (*4*).

*Brain-state analysis in the primary cohort*. In the primary cohort, brain-state features were defined at the group level and brain-state scores were computed for each individual across learning stages. For each participant and scan, preprocessed fMRI time series were extracted from a whole-brain cortical parcellation comprising 400 cortical regions. Regional activity variability was quantified as the temporal standard deviation of each regional time series. Regions exhibiting significantly higher variability in the learning group compared with controls were identified using one-sided independent-samples $t$-tests ($P < 0.05$), defining a set of fluctuation-changed regions. We next assessed whether these regions also exhibited altered coordination. Subject-level pairwise functional coupling (absolute Pearson's correlation) was computed among all fluctuation-changed regions. Region pairs showing significantly higher coupling in the learning group relative to controls were identified using one-sided independent-samples $t$-tests ($P < 0.05$) and retained, yielding a subnetwork jointly characterized by coordinated changes in regional variability and inter-regional functional coupling. To summarize the expression of this brain state at the individual level, we computed a brain-state score for each participant as the product of the average magnitude of regional fluctuations and the average strength of functional coupling within the identified subnetwork. Higher scores indicate a greater departure from steady-state organization toward a coordinated, fluctuation-altered non-stationary brain state.

*Brain-state analysis in the validation cohort*. In the validation cohort, the same brain-state analysis framework was applied at the individual level to track longitudinal brain-state dynamics under dense sampling. Given the within-participant longitudinal design, no group-level statistical tests were performed. Instead, brain-state scores were computed longitudinally across learning stages within participants to examine whether the dynamical patterns identified in the primary cohort emerged at the individual level independently. Dataset-specific adaptations were implemented to accommodate differences in study design while preserving the conceptual definition of coordinated variability–coupling changes.

Network analysis

To further characterize learning-related reorganization of functional brain architecture, we conducted network analyses within each participant across stages. For each participant and scan, preprocessed fMRI time series were extracted from the same 400-region whole-brain cortical parcellation (*5*, *6*). Functional connectivity matrices were constructed by computing pairwise Pearson's correlations between regional time series.

*Modularity analysis.* Community structure was identified using repeated Louvain partitioning to ensure robust detection of modular organization. Across multiple iterations, consensus community assignments were derived, and the consistency of regional module affiliation was quantified. To characterize learning-related changes in network organization, recruitment and integration metrics were computed (*7*). Recruitment quantified the extent to which regions preferentially interacted within their assigned network, reflecting within-network cohesion. Integration quantified the degree of interaction between different networks, capturing cross-network communication. These metrics were computed at each stage for each participant, enabling assessment of how learning reshaped the brain network segregation and integration.

*Network connections analysis*. To ensure comparability of network connections across participants and stages, proportional thresholding was applied to each functional connectivity matrix, retaining the strongest 5% of connections. This approach controlled for inter-individual differences in overall connectivity strength while preserving the most robust functional links. Following thresholding, the number of intra-network and inter-network connections was quantified for each participant and stage. These measures were used to assess how learning selectively modulated connectivity within and between functional networks over time.

Statistical analysis

*Linear mixed-effects (LME) model*. Longitudinal changes in behavioral and neuroimaging measures were assessed using LME models to account for repeated measurements within individuals. All models included subject-level random intercepts and fixed effects of group, learning stage, and sex as a covariate.

To evaluate overall learning-related effects, a baseline model included main effects of group and stage. To explicitly test whether learning trajectories differed between groups, an interaction model additionally included the group × stage interaction. Model formulations were as follows:

model 1: outcome ~ group + stage + sex + (1 | subject)

model 2: outcome ~ group + stage + group × stage + sex + (1 | subject)

Statistical significance of interaction effects was assessed using standard inferential procedures for mixed-effects models. When relevant, post hoc contrasts were performed to characterize stage-specific group differences.

*Feature importance estimation*. To evaluate multivariate separability of behavioral profiles between groups, a Random Forest classifier was trained on cognitive measures across learning stages. Model performance was assessed using five-fold cross-validation, and classification accuracy, F1-score, and area under the receiver operating characteristic curve (AUC) were reported. Feature importance estimates were extracted to quantify the relative contribution of each cognitive metric to group discrimination.

### Supplementary Text

AMO learning protocol

AMO learning involves progressive transition from physical abacus manipulation to fully internalized mental arithmetic. It begins with explicit motor-guided computation and gradually shifts toward motor-independent mental visualization of the abacus. With increasing expertise, reliance on finger movements is progressively reduced, and participants perform increasingly complex arithmetic operations using internalized representations. This staged design enables investigation of learning-dependent transformation from embodied to abstract computation.

Efficacy of abacus learning

We assessed AMO proficiency in children enrolled in the primary dataset using the standardized AMO level test, in which Level 1 denotes the highest performance tier, requiring mental arithmetic with 10 operands of 2–4 digits. The assessment was conducted under time constraints and required participants to complete all items using AMO strategies. Learning efficacy was evaluated by comparing proficiency levels at the late learning stage. Among the assessed trainees, 93% achieved high proficiency, with 87% reaching Level 1 and an additional 4% attaining Level 2, indicating robust acquisition of AMO skills following learning.

Cognitive assessment battery of primary cohort

Cognitive outcomes were assessed using standardized measures of mathematical ability and executive function, including arithmetic computation, visuospatial numerical processing, numerical magnitude representation, inhibitory control, and cognitive flexibility.

*Heidelberg Rechentest.* The HRT yields two subscale scores (*8*): arithmetic ability, assessed by six timed subtests (addition, subtraction, multiplication, division, equation completion, and numerical comparison), and visuospatial mathematical ability, assessed by five timed subtests (line estimation, pictorial enumeration, three-dimensions cube counting, sequential number counting, and numerical pattern sequencing). A number-copying subtest preceded formal testing to familiarize participants with task demands. Raw scores were converted to age-normed T scores based on Chinese urban population norms. Composite scores for arithmetic and visuospatial abilities were computed by averaging T scores within each subscale, yielding two summary metrics indexing arithmetic and visuospatial mathematical ability.

*Compare-to-5 task.* Participants classified single digits (1–9, excluding 5) as larger or smaller than 5. Digits were grouped by numerical distance into proximal (3, 4, 6, 7) and distal (1, 2, 8, 9) conditions. The task comprised 96 trials (48 per condition), presented in a randomized order with no consecutive repetitions. Each trial began with a 500ms fixation, followed by the target stimulus presented until response, and ended with a 500ms fixation. Mean reaction time for correct trials and response accuracy were computed separately for each condition. Reaction time and accuracy costs between proximal and distal conditions were then calculated and used as indices of numerical magnitude processing efficiency.

*Go/No-go task.* Participants responded to pictures of animals (go trials) and withheld responses to pictures of chimpanzees (no-go trials). Stimuli were displayed for 500 ms with an inter-stimulus interval of 1100–1200 ms. After 12 practice trials, two experimental blocks of 70 trials each were administered; no-go trials comprised 20% of trials to establish a prepotent response tendency. The inhibition cost was quantified by normalizing the reaction time on correct go trials by go-trial accuracy across all trials, serving as an index of inhibitory efficiency.

*Dots task.* This task assessed cognitive flexibility across three blocks. In the congruent block, participants responded ipsilaterally to one dot type (striped or grey), whereas in the incongruent block, they responded contralaterally to the other dot type. The mixed block (61 trials) intermixed both stimulus–response mappings, requiring participants to switch between rules. Each trial consisted of a 500ms fixation, a 500ms interstimulus interval, target presentation (up to 750ms or until response), and a 500ms post-response interval. Stimulus–response mappings were counterbalanced across participants. For each condition, mean reaction time for correct trials and response accuracy were computed as indices of shifting ability. Switching cost was calculated as the difference in reaction time and accuracy between switch and non-switch trials within the mixed block.

MRI acquisition and preprocessing

*MRI acquisition and preprocessing of primary longitudinal dataset*. Resting-state fMRI data were acquired in 2009 using a 1.5 T Philips Achieva scanner with an eight-channel head coil. A total of 180 volumes during resting state were collected over 360s with a single-shot EPI sequence (TR = 2,000ms, TE = 50ms, flip angle = 90°, slice thickness = 5mm, slice gap = 0.8mm, FOV = 230 × 230mm², matrix = 64 × 64, 22 interleaved ascending slices). High-resolution T1-weighted images were obtained using a 3D fast field-echo sequence (TR = 25ms, TE = 4.6ms, flip angle = 15°, FOV = 256 × 256mm², matrix = 256 × 256, voxel size = 1 × 1 × 1mm³, 150 sagittal slices). Preprocessing was carried out with DPARSF-A (v4.2). The first five volumes were discarded for signal stabilization. Remaining volumes underwent slice-timing correction and motion realignment. Nuisance regressors (white matter, cerebrospinal fluid, global signal, and 24 head-motion parameters) were regressed out. Functional images were normalized to MNI space (resampled to 4 mm isotropic voxels), smoothed with a 6-mm FWHM Gaussian kernel, linearly detrended, and band-pass filtered (0.01–0.1 Hz). Region-of-interest time series were then extracted for subsequent connectivity analyses.

*MRI acquisition and preprocessing of dense-sampling validation dataset*. MRI scans were acquired on a 3T Siemens Prisma scanner using a 20-channel head coil. Rs-fMRI scans were collected with a multi-echo, multiband EPI sequence (TR = 1,500ms; TE = 12.60/30.86/49.12/67.38ms; FA = 73°; multiband factor = 3; voxel size = 3mm isotropic; 48 slices; FOV = 216 × 216mm²). Two resting-state runs (280 volumes, 7min each) were acquired per visit with eyes open and central fixation. High-resolution T1-weighted images were acquired using a 3D MPRAGE sequence (TR = 2,300ms; TE = 2.32ms; TI = 900ms; FA = 8°; voxel size = 0.7mm isotropic). Rs-fMRI scans were preprocessed using an in-house pipeline optimized for multi-echo acquisitions. Preprocessing included de-spiking, slice-timing correction, and motion correction. Multi-echo denoising was performed using TEDANA, with T2*-weighted optimal echo combination followed by multi-echo ICA (ME-ICA) to classify and remove non-BOLD (S0-dependent) components. Data were nonlinearly registered to MNI space, projected onto fs_LR 32k mid-thickness surfaces, and combined with subcortical data in CIFTI format. Spatial smoothing was applied using Gaussian kernels (σ = 2.55mm). Nuisance regression included white matter, cerebrospinal fluid, and global signals. Data were band-pass filtered (0.01–0.1 Hz), and volumes with framewise displacement > 0.3mm were censored.

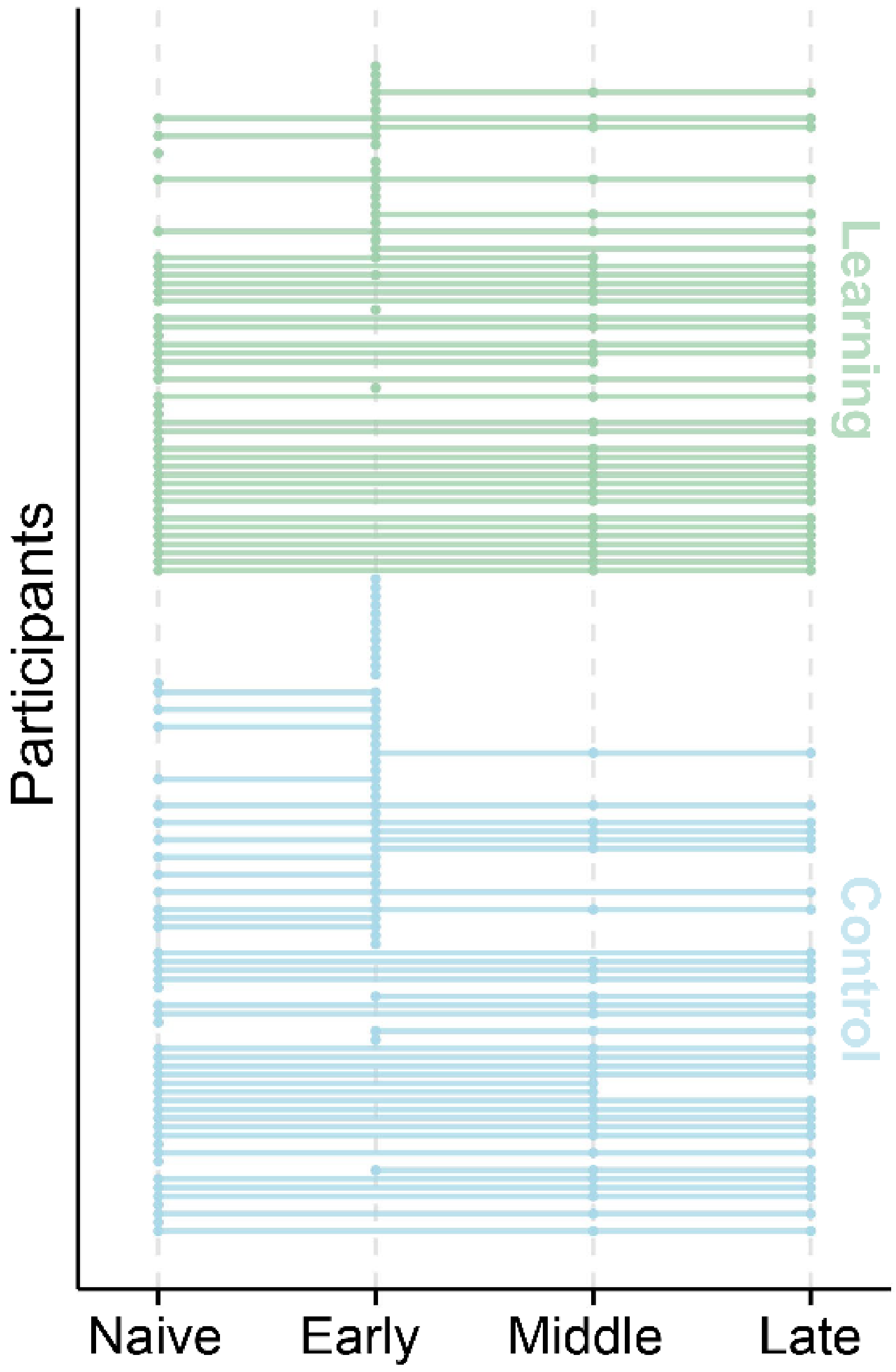


**Fig. S1.** Rs-fMRI visits of primary longitudinal dataset.

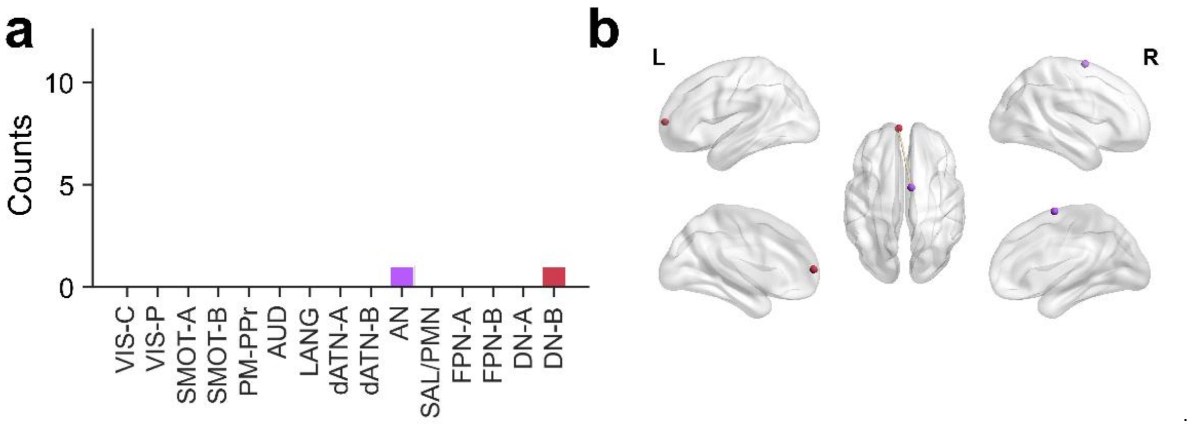


**Fig. S2.** There existed no significant difference of brain network between groups in naive stage.

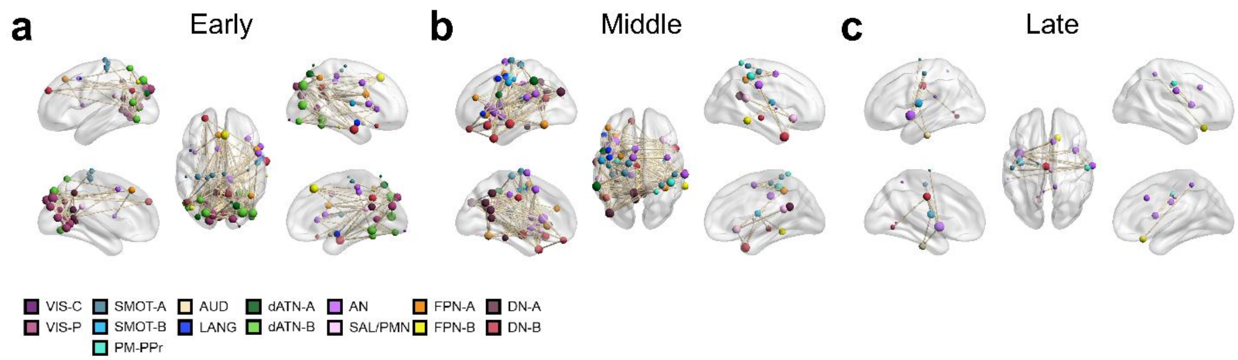


**Fig. S3.** Brain networks contributing to the significant group difference of brain-state scores during learning stages.

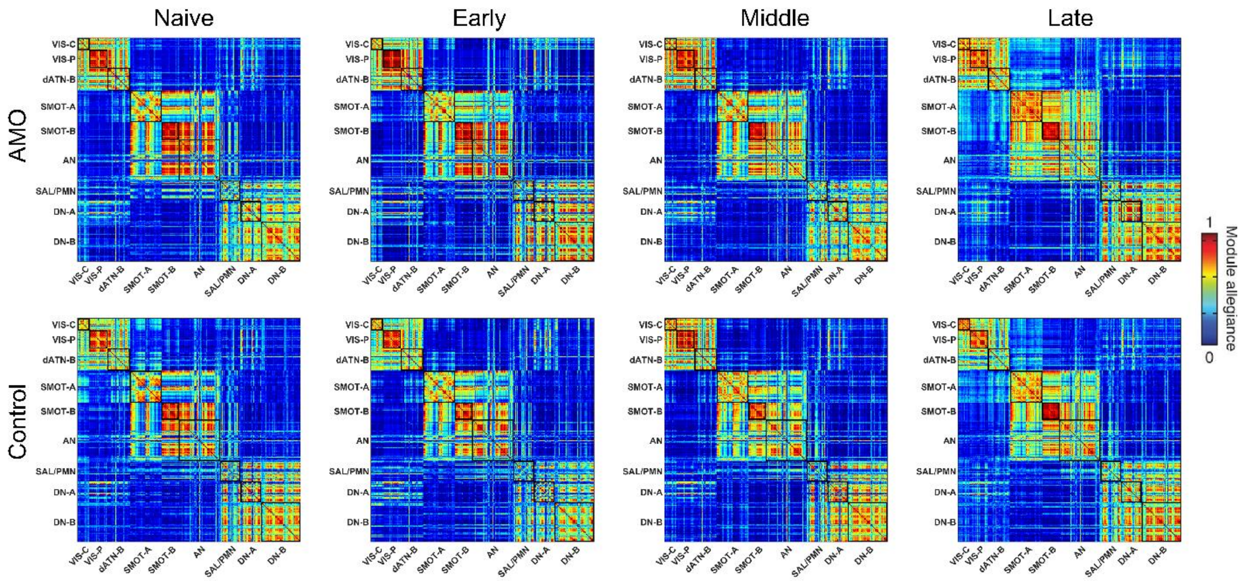


**Fig. S4.** Module allegiance matrices across learning stages.

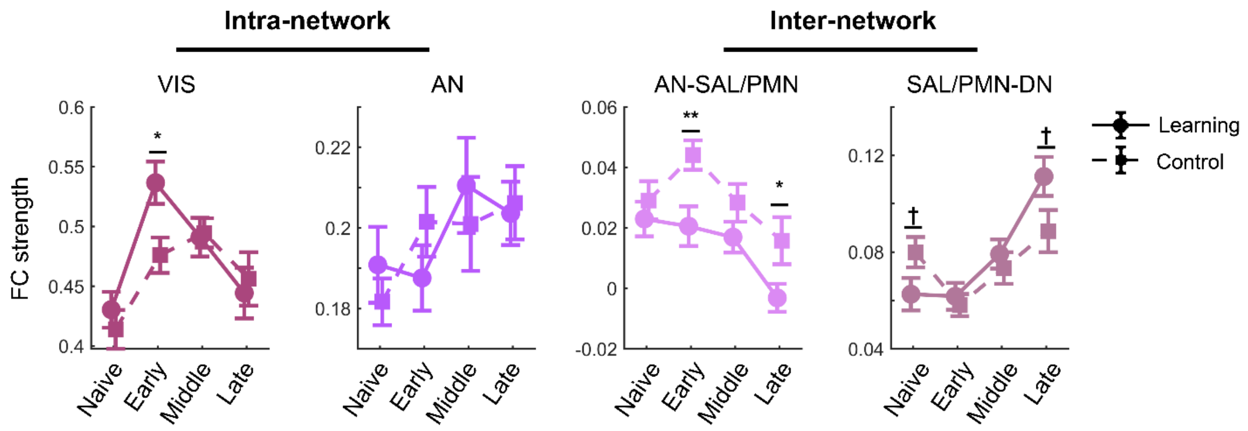


**Fig. S5.** Reweighting of intra- and inter-network functional connectivity strength across learning stages. Note: ***$P < 0.001$, **$P < 0.01$, *$P < 0.05$, †$P > 0.05$.

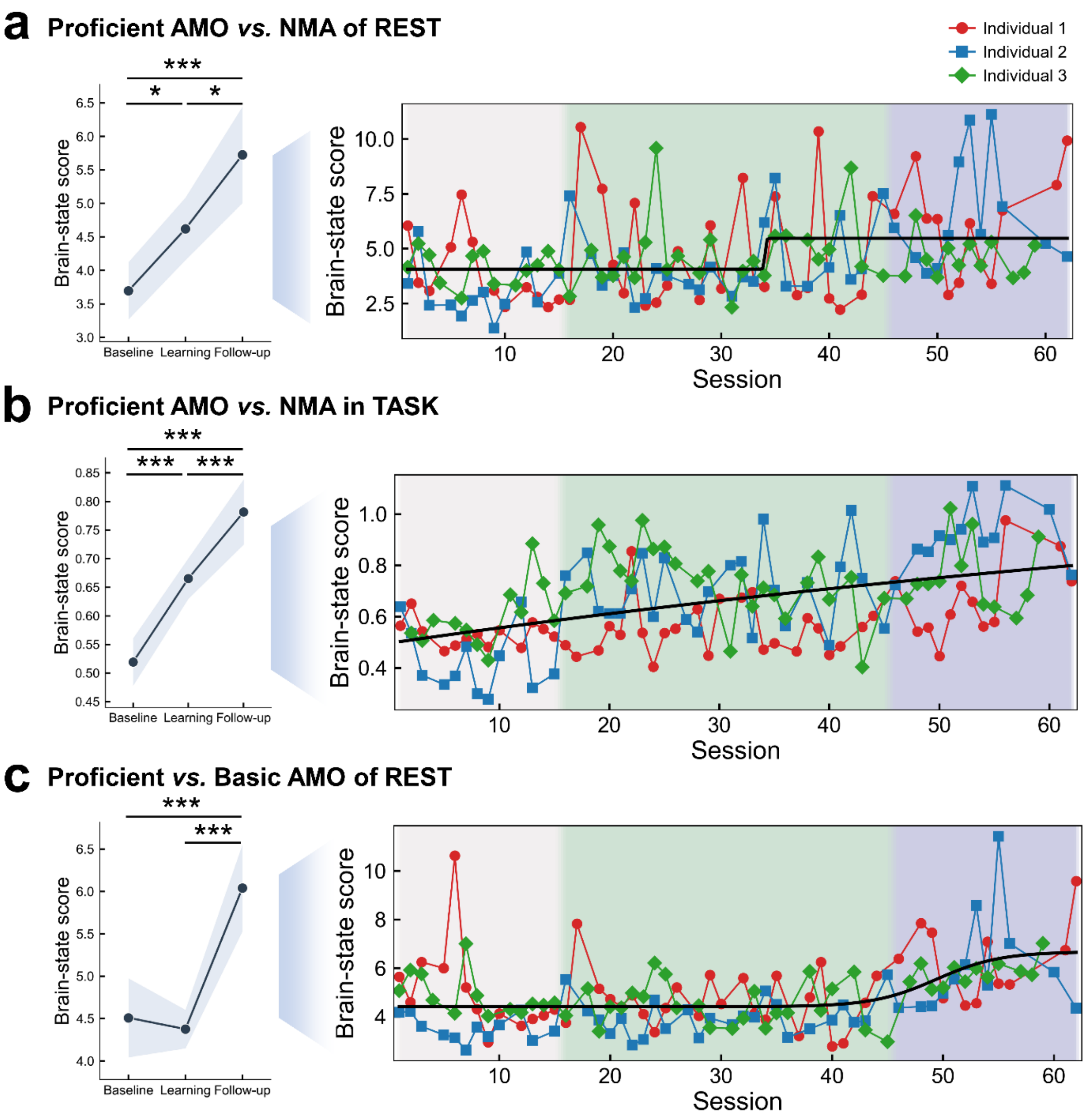


**Fig. S6.** Brain-state scores across stages and individual trajectories with independent validation dataset. Note: ***$P < 0.001$, *$P < 0.05$.

**Table S1.** Demographics of primary longitudinal cohort.

| Demographics | Group | | Statistics |
|---|---|---|---|
| | Learning | Control | |
| age (mean ± SD) | 6.89 ± 0.53 | 6.95 ± 0.52 | $t(184) = -0.865$, $p = 0.388$ |
| sex (total, female) | 87 (48) | 99 (48) | $\chi^2 = 0.829$, $p = 0.362$ |

**Table S2.** Baseline control of cognitive variables in primary longitudinal cohort.

| Variables | | Group | | Statistics |
|---|---|---|---|---|
| | | Learning | Control | |
| IQ | Raven scores | 102 ± 14 | 102 ± 12 | *t*(159) = -0.254, *p* = 0.800 |
| Early school behavior rating scale | conduct | 14.8 ± 3.3 | 14.6 ± 3.0 | *t*(160) = 0.338, *p* = 0.736 |
| | internal | 31.2 ± 5.2 | 30.0 ± 5.4 | *t*(160) = 1.486, *p* = 0.139 |
| | competence | 42.6 ± 6.4 | 43.0 ± 7.6 | *t*(160) = -0.354, *p* = 0.724 |
| Mastery motivation | object-oriented persistence | 3.32 ± 0.55 | 3.33 ± 0.61 | *t*(156) = -0.126, *p* = 0.900 |
| | social persistence with adults | 3.81 ± 0.68 | 3.94 ± 0.62 | *t*(156) = -1.255, *p* = 0.211 |
| | social persistence with children | 4.00 ± 0.65 | 4.14 ± 0.57 | *t*(156) = -1.445, *p* = 0.150 |
| | gross motor persistence | 3.58 ± 0.62 | 3.64 ± 0.61 | *t*(156) = -0.701, *p* = 0.485 |
| | mastery pleasure | 3.94 ± 0.45 | 4.04 ± 0.56 | *t*(156) = -1.314, *p* = 0.191 |
| | negative reaction to failure | 3.32 ± 0.66 | 3.42 ± 0.61 | *t*(156) = -1.023, *p* = 0.308 |
| | general competence | 3.43 ± 0.65 | 3.48 ± 0.64 | *t*(156) = -0.563, *p* = 0.574 |
